# Disruption of Tonic Endocannabinoid Signaling Triggers the Generation of a Stress Response

**DOI:** 10.1101/2022.09.27.509585

**Authors:** Gavin N. Petrie, Georgia Balsevich, Tamás Füzesi, Robert J. Aukema, Wouter P. F. Driever, Mario van der Stelt, Jaideep S. Bainsand, Matthew N. Hill

## Abstract

Endocannabinoid (eCB) signalling gates many aspects of the stress response, including the hypothalamic-pituitary-adrenal (HPA) axis. The HPA axis is controlled by corticotropin releasing hormone (CRH) producing neurons in the paraventricular nucleus of the hypothalamus (PVN). Disruption of eCB signalling increases drive to the HPA axis, but the mechanisms subserving this process are poorly understood. Using an array of cellular, endocrine and behavioral readouts associated with activation of CRH neurons in the PVN, we evaluated the contributions of tonic eCB signaling to the generation of a stress response. The CB1 receptor antagonist/inverse agonist AM251, neutral antagonist NESS243, and NAPE PLD inhibitor LEI401 all uniformly increased c-fos in the PVN, unmasked stress-linked behaviors, such as grooming, and increased circulating CORT, recapitulating the effects of stress. Similar effects were also seen after direct administration of AM251 into the PVN, while optogenetic inhibition of PVN CRH neurons ameliorated stress-like behavioral changes produced by disruption of eCB signaling. These data indicate that under resting conditions, constitutive eCB signaling restricts activation of the HPA axis through local regulation of CRH neurons in the PVN.

## Introduction

Exposure to a threat or challenge elicits a host of physiological and psychological consequences collectively known as the stress response. Although the body’s reaction to stress can be beneficial in acute conditions, unmitigated activation of stress-responsive biological systems can lead to negative psychological, metabolic, and cellular consequences (McEwen et al., 2015). While the neural mechanisms which regulate the generation of a stress response following exposure to a challenge have been relatively well characterized, the process by which stress-responsive systems are constrained during ambient conditions is less well understood (Herman et al., 2016). As many disease states, such as mood and anxiety disorders as well as many inflammatory conditions, are associated with aberrant activation of stress-responsive systems in the absence of overt threats or challenges, understanding the neural processes involved in preventing undue activation of stress-responses is an important area that has been historically neglected.

There are multiple, well-characterized arms of the stress response; amongst these, the neuroendocrine response via hypothalamic-pituitary-adrenal axis (HPA) activation is the most well understood (Pecoraro et al., 2006). Activation of the HPA axis begins with the release of corticotropin releasing hormone (CRH) from parvocellular neurons in the paraventricular nucleus of the hypothalamus (PVN) and ends with elevated circulating corticosterone (CORT) (Herman et al., 2010). Alongside HPA activation, stress exposure triggers a complex behavioral response in animals including alterations in vigilance, avoidance and threat assessment. While these behavioral responses have traditionally been measured in a spurious manner, recent work has described a reliable stress-induced pattern of behavioral changes, with emphasis on an increase in stereotypic grooming (Füzesi et al., 2016; Loewen et al., 2020).

The endocannabinoid system (eCB) is intimately involved with the stress response and has been widely studied as a system which dynamically responds to stressful experiences and modulates synaptic plasticity across multiple brain circuits to influence shifts in behavioral repertoires (Morena et al., 2016). The two primary eCB ligands anandamide (AEA; Devane et al., 1992) and 2-arachidonoyl glycerol (2-AG; Mechoulam et al., 1995; Sugiura et al., 1995) are primarily synthesized postsynaptically by their respective synthetic enzymes n-acylphosphatidylethanolamine phospholipase D (NAPE-PLD; Cadas et al., 1997; Okamoto et al., 2004) and Diacylglycerol lipase (DAGL; Sugiura et al., 1995; Bisogno et al., 2003) and then act retrogradely on presynaptic cannabinoid receptors (CB1; Matsuda et al., 1990; Wilson et al., 2001) to dampen synaptic transmitter release. Both AEA and 2-AG have been implicated in the regulation of the HPA axis, particularly in the context of termination and recovery from stress exposure by modulating excitability in stress-responsive neural circuits, through acting at the CB1 receptor (Morena et al., 2016). In addition to this dynamic role of eCB signaling in response to stress exposure, there is also some evidence that eCB signaling may also play a role in tonic suppression of the stress response in ambient conditions. For example, AEA signaling is known to rapidly decline in response to stress, and inhibition of its metabolism can blunt many aspects of stress suggesting that tonic AEA signaling may be constraining activation of the stress response in the absence of a threat (Bedse et al., 2017, 2014; Bluett et al., 2014; Hill et al., 2009; Gray et al., 2015; Sticht et al., 2019; Yasmin et al., 2020), while depletion of AEA can increase CORT secretion (Mock et al., 2020). Consistent with this, administration of a CB1 receptor antagonist reliably increases circulating glucocorticoids, suggesting that disruption of tonic eCB signaling is sufficient to increase activation of the HPA axis (Hill et al., 2009; Newsom et al., 2020, 2012; Patel et al., 2004; Steiner et al., 2008; Wade et al., 2006). While the neural circuits through which dynamic changes in eCB signaling regulate stress responses is somewhat characterized (Morena et al., 2016), there is a relative paucity of information regarding the mechanism and circuits by which tonic eCB signaling maintains quiescence of the HPA axis under ambient conditions. Using an array of anatomical, neuroendocrine and behavioral readouts, we combine multiple pharmacological approaches to disrupt tonic eCB signaling to elucidate the nature by which tonic eCB signaling constrains stress responsive processes.

## Methods

### Animals

All experiments were approved by the University of Calgary Animal Care and Use Committee in accordance with Canadian Council on Animal Care Guidelines. Transgenic Crh-IRES-Cre (B6(Cg)-Crhtm1(cre)Zjh/J; stock number 012704)/Ail4 (B6.Cg-Gt(ROSA)26Sortm14(CAG-TdTomato0Hze/J; stock number 007914) mice, who have been previously described in (Cusulin et al., 2013; Madisen et al., 2010; Taniguchi et al., 2011), were bred in house to express td-tomato in CRH expressing neurons. Wild type (C57BL/6J; stock number 000664) mice were purchased from Jackson laboratories. For viral and CRH neuron activity experiments, offspring derived from crosses of homozygous Crh-IRES-Cre and Ai14 genotypes were used. Mice were single housed on a 12:12h light dark schedule where lights were turned on at 8:00 am. Before testing at 10-12 weeks old, animals were housed in Greenline ventilated cages containing small wood chip bedding, shredded paper and a paper or plastic house for environmental enrichment.

### Drugs

All systemically administered drugs were dissolved in 5% polyethylene glycol, 5% Tween- 80, and 90% saline the day of each experiment. The CB1 receptor antagonist/inverse agonist AM251 (3mg/kg; Cayman Chemical, Ann Arbor, MI, USA) and the neutral CB1 receptor antagonist NESS0327 (0.3mg/kg; Cayman Chemical, Ann Arbor, MI, USA) were administered 15 minutes prior to behavioral recording. The NAPE-PLD inhibitor LEI401 (30mg/kg; was synthesized as previously described; Mock et al., 2020) and the DAGL inhibitor DO34 (30mg/kg; was synthesized as previously described; Ogasawara et al., 2016) were administered I.P. 120 minutes before behavioral recording. For central administrations, AM251 (Cayman Chemical, Ann Arbor, MI, USA) was mixed in 80% DMSO and 20% saline to make a 1ug/200nl solution and 200nl was administered unilaterally into the PVN, or bilaterally into the striatum, 15 minutes before behavioral recording. Drug was infused through 30-gauge injection needle connected to a 10ul Hamilton micro syringe by polyethylene tubing and driven by a minipump (Harvard Apparatus, Holliston, MA, USA). A three-minute infusion protocol consisted of one-minute active infusion and a two-minute diffusion period to allow for proper spread and prevent back flow into the cannula.

### Behavioral protocol

All animals were handled for four days where they were familiarized with the experimenter and drug administration procedures. After handling, animals were habituated an hour a day for three days to their testing room and “home cage” environment, which consisted of wood chip bedding, while their shredded paper, house, food hopper and water bottle were removed. On test day, animals had their enrichment and food/water removed an hour before behavioral measurements. Mice then received drug, or vehicle, administrations before behavioral assessment. Following drug administration, behaviors were recorded in their home cage for a 15- minute epoch.

Recorded behavior videos were scored using a custom python behavioral scoring program by an individual blinded to each subject’s treatment group. Seven behaviors were scored (as described by Füzesi et al., 2016) as time spent: walking (movement of front and back paws to change location or direction); surveying (sniffing of environment while back legs remained stationary); rearing (achieving an upright position by lifting of front legs and simultaneous extension of back legs); digging (displacing bedding); chewing (chewing of any inanimate object in the cage); grooming (body licking, rubbing or scratching); resting (no observable movement).

### Immunohistochemistry

On each test day, 90 minutes after behavioral monitoring began, animals were deeply anaesthetized using pentobarbital and perfused intracardially with 4% paraformaldehyde (PFA) and phosphate buffered saline (PBS). Following perfusion, the brains were extracted and post-fixed in 4% PFA at 4 ° C overnight and then submerged in 30% sucrose in PBS at 4 ° C for 48 hours. Brains were flash frozen, mounted and sectioned (40µm) on a Leica SM 2010R sliding microtome. Slices containing the PVN were collected according to Paxinos and Franklin’s mouse atlas and placed in an anti-freeze solution until antibody treatment.

Sections were washed three times (all washes are 10 minutes in duration) in a PBS solution and then three more times in a PBS and 0.1% Triton X-100 solution (PBST) before incubation in PBS blocking buffer containing 5% normal donkey serum at room temperature for 1h. Sections were then incubated in primary antibody (rabbit anti-c-Fos; 1:300 Cell Signaling Technology, Danvers, MA, USA) at 4 ° C overnight. Tissue was then washed in PBST solution three times and then incubated in secondary antibody at room temperature for two hours (donkey anti-rabbit Alexa Fluor 488 [1:100 Jackson Laboratory, Farmington, CT, USA] or Alexa Fluor 647 donkey anti rabbit [1:100 J Jackson Laboratory, Farmington, CT, USA] for colocalization studies). Sections were then mounted in PBS onto glass slides using Fluoroshield with DAPI (Sigma-Aldrich) and examined using a Leica TCS SPE-II confocal microscope (confocal is Leica TCS SPE-II and a fluorescent is Leica DM4000 B/M LED). Two or three slices containing the PVN were collected from each specimen. Gray-scale images were taken of both hemispheres separately for each slice using I3 and N21 filtercubes. All images from the same experiment were obtained under the same exposure and magnification (20X; NA 0.55). Images were processed in ImageJ, where they were subjected to an identical brightness threshold before c-fos labeled cells were counted based on size criteria set for each experiment. A boundary of the entire PVN was drawn according to the Paxinos and Franklin’s mouse atlas and the total number of FOS+ cells in this area were counted. This was then normalized to the size of the PVN for each slice by dividing number of FOS+ cells by area, to give a cells/mm^2^ value that could be compared across drug groups. For colocalization studies, slices were imaged using a Leica TCS SPE-II confocal microscope. Cells expressing both td-tomato (CRH expressing neurons) and c-fos were identified in the manner described above and z stacks were manually assessed on the microscope to confirm colocalization. Number of c-fos+/td-tomato+ co-expressing cells were divided by the total number of td-expressing cells within the same region to obtain a percent value for each animal.

For histology, perfusions and slicing were performed in the same way as described for immunohistochemistry. Brain slices were mounted and stained for DAPI to identify cell bodies. Slices were then imaged using a fluorescent microscope using the DAF filter cube to visualize DAPI

### Corticosterone Analysis

Tail blood was taken 15 minutes following behavioral recordings using microvette collection tubes and stored in wet ice until spinning. Blood samples were centrifuged at 10,000 x g for 5 min at 20°C. Serum was stored at -20°C until analysis in triplicates with a Corticosterone ELISA kit (Arbor Assays, Ann Arbor, MI, USA) by following the manufacturer’s instructions.

### Injection and implantation

An automatic (Neurostar) stereotaxic apparatus (Kopf Instruments) outfitted with an isoflurane delivery setup was used for all surgical procedures. Mice were placed under deep anesthesia via isoflurane prior to being securely placed into ear bars. All animals received subcutaneous injections of saline (1ml) and Metacam (2mg/kg). The scalp was shaved and 30% ethanol was applied prior to incision. The skull was exposed and holes were drilled to allow for implantation of cannulae or injection. For drug infusion experiments, a unilateral cannula was lowered into the PVN (anteroposterior (AP), -0.25 mm; lateral (L), 0.3 mm from bregma; dorsoventral (DV), -4.5 mm from dura) and secured with Metabond and dental cement. For optogenetic studies, a pulled glass capillary was lowered into the PVN (AP, -0.3 mm; L, 0.2 mm from bregma; DV, -4.8 mm from dura) and 210nl of Arch 3.0- eYFP (rAAV2-EF1a-double floxed-eArch3.0-eYFP; 5 × 10^11^ GC per ml; UNC Vector Core or eYFP (Addgene plasmid 20296, pAAV-EF1a-double floxed-eYFP-WPRE-HGHpA; 5 × 10^11^ GC per ml; Penn Vector Core, Philadelphia, PA, USA) was infused using a Drummond Nanoject III in three 70nl bolouses. Mice were sutured and allowed to recover. Two weeks later, mice had 400µm diameter optical ferrules (400/460/0.48; Doric, Franquet, QC, CA) lowered over the PVN (AP, -0.3 mm; L, 0 mm from bregma; DV, -4.5 mm from dura) and secured similarly to cannula implants. All animals were given at least 1 week to recover following surgical procedures before the initiation of behavioral experiments.

### Optogenetic procedures

A light source (593 nm, IKE-593-100-OP, IkeCool Corporation) was connected to the implanted ferrule with a 2-meter fibre optic cable (400/460/1100-0.48; Doric, Franquet, QC, CA). Animals underwent four handling and three habituation days prior to behavioral testing. During habituation, mice were attached to the fiber and allowed to wander their home cage for an hour. AM251 (3mg/kg) or VEH (10µl/g) injections were administered i.p. 15 minutes prior to a 15-minute behavioral recording during which all animals underwent exposure to continuous yellow light (532nm;15mW). Following behavioral testing, mice were perfused and PVN CRH neurons were assessed for viral expression using a Leica DM4000 B/M LED fluorescent microscope at 10x magnification.

## Results

### Disruption of eCB signaling activates a stress-like response

To investigate the effects of CB1 receptor blockade on the stress response, we administered the CB1 receptor antagonist/inverse agonist AM251 (3mg/kg, *i*.*p*.) before probing three well established arms of the stress response: 1) c-fos induction of CRH expressing neurons in the PVN; 2) elevations in circulating CORT; 3) increases in repetitive behaviors (e.g., grooming) in a homecage environment (Fig.1a). When compared to an intraperitoneal (I.P.) vehicle administration, AM251 (I.P.) significantly altered the homecage behavioral profile of mice, as shown by the raster plots depicting the behaviors of each mouse over the 15-minute epoch (Fig.1b). Specifically, AM251 administered animals spent significantly more time walking t(14) = 3.892, p = 0.0016, rearing t(14) = 4.291, p = 0.0007, and grooming t(14) = 2.153, p = 0.0492, paralleling behavioral changes in mice exposed to acute stress (Füzesi et al., 2016; Fig.1c). In further support of disruption of eCB signaling producing a stress-like response, CORT levels were measured in the blood 30 minutes following behavior recording onset. AM251 significantly increased circulating CORT levels when compared VEH t(12) = 3.525, p = 0.0042 (Fig. 1d). AM251 also increased c-fos expression in the PVN when compared to VEH t(13) = 2.950, p = 0.0113. Furthermore, focusing in on the stress-integrative PVN CRH neurons, AM251 administration dramatically increased c-fos expression in these neurons t(14) = 5.769, *p = <0*.*0001* (Fig. 1e, f). Together, these results highlight the importance of eCB signaling in constraining activation of a stress-like response in a low stress, familiar environment.

**Figure 1.**
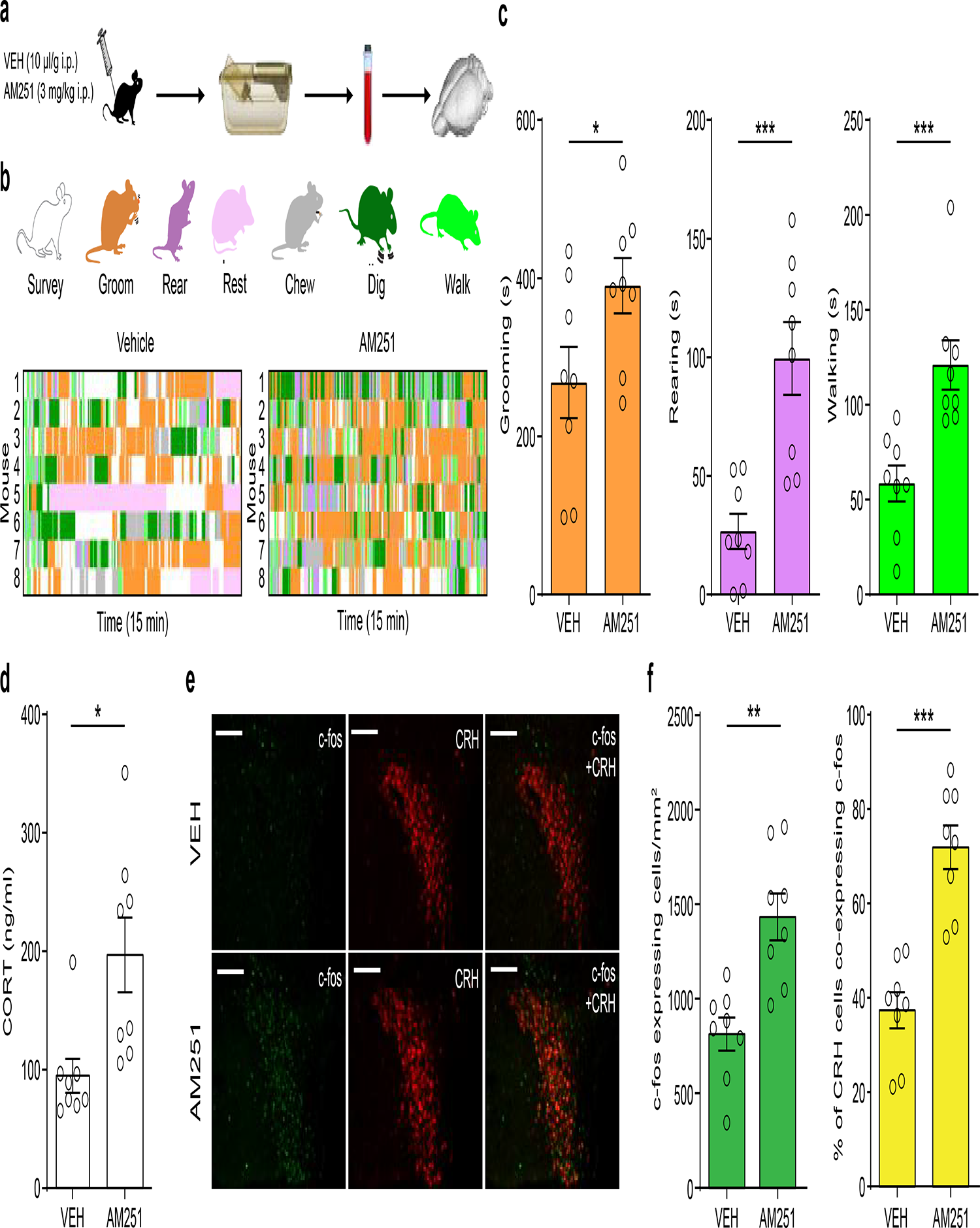
CB1 blockade precipitates a stress-like phenotype. (a) Procedural outline showing drug administration 15 minutes before home cage behavioral recording, 45 minutes before blood collection and 105 minutes before brain extraction. (b) Raster plot shows the detailed behavioural output throughout the 15-minute epoch, where each behavior is color coded based on the provided index. (c) Quantification of grooming (t(14) = 2.153, p = 0.0492), rearing (t(14) = 4.291, p = 0.0007) and walking (t(14) = 3.892, p = 0.0016) behaviors following VEH or AM251. (d) Corticosterone (CORT) levels following Vehicle (VEH) or AM251 administration (t(12) = 3.525, p = 0.0042) (e) c-fos immunohistochemical activity (green) in CRH expressing neurons (red) in the PVN following VEH (left) or AM251 (right). (f) Quantification of c-fos expressing cells (Left; t(13) = 2.950, p = 0.0113) and % of CRH expressing cells that co-expressed c-fos (right; t(14) = 5.769, p =, 0.0001). Scale bars: 150um *p = <0.05 **p = <0.01 ***p = <0.001.

### Pharmacological characterization of tonic eCB regulation of the stress-like response

The CB1 receptor is a G protein-coupled receptor with a high degree of constitutive signaling: tonic eCB signaling can be both agonist-dependent (mediated by the release of AEA or 2-AG) or agonist-independent (constitutive G protein signaling). To establish if tonic eCB regulation of the stress-like response is driven by eCB signaling itself or via constitutive receptor activity, we employed the use of the neutral CB1 receptor antagonist NESS0327 (I.P.; Fig.1a), which, unlike AM251, blocks ligand dependent activation of CB1 receptors but does not prevent agonist-independent constitutive signaling (which is blocked by AM251). In comparison to VEH, mice administered NESS showed significantly more homecage grooming behaviors t(13) = 3.089, p = 0.0086 (Fig. 2b), elevated circulating CORT levels t(13) = 2.318, p = 0.0374 (Fig. 2c), and elevated c-fos expression in PVN neurons t(13) = 2.459, p = 0.0287 (Fig. 2d,e). These results suggest that active, tonic eCB signaling of AEA and/or 2-AG at CB1 receptors governs inhibition of stress responsive networks at rest.

**Figure 2.**
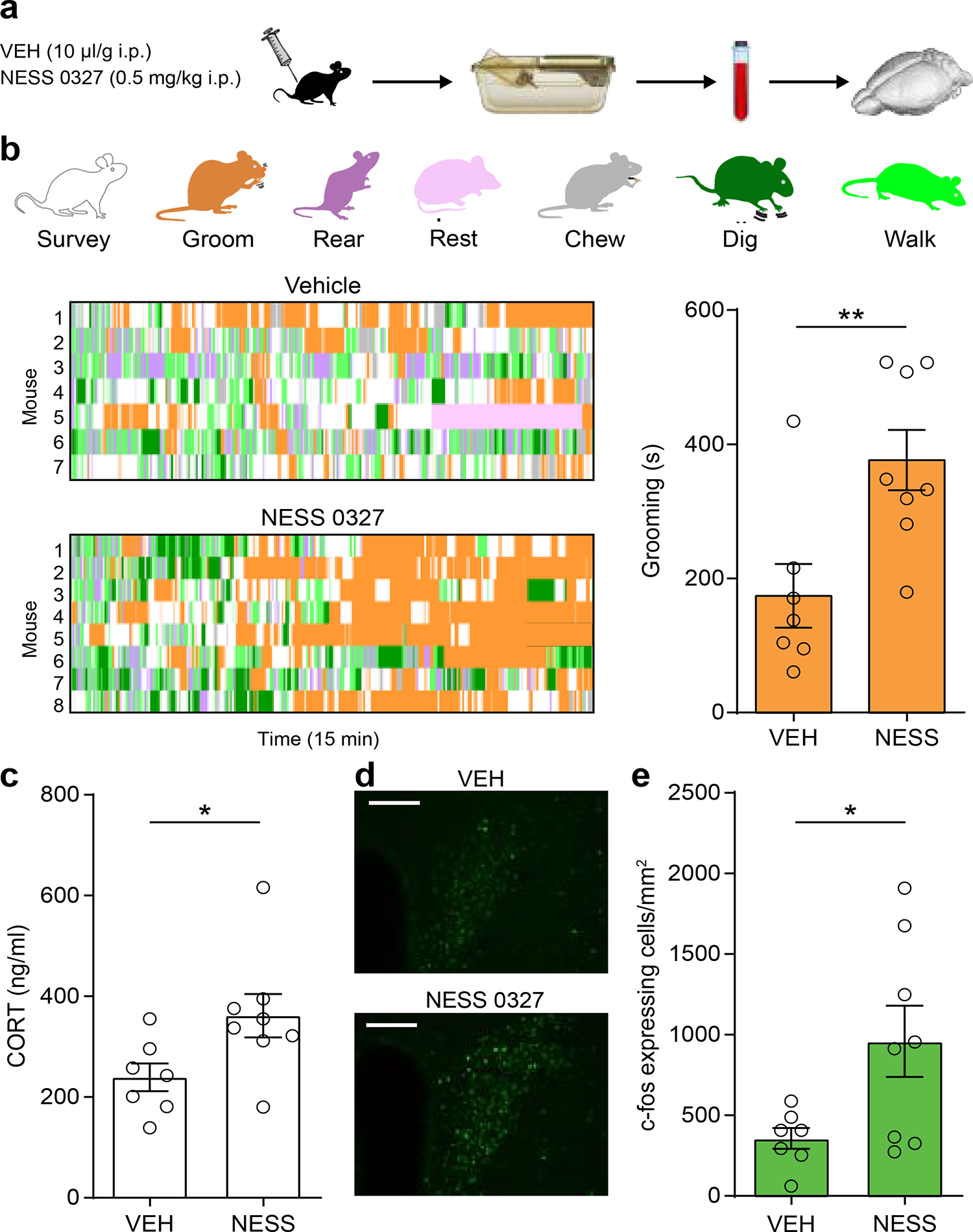
Neutral CB1 antagonist initiates the stress response. (a) Mice were administered NESS or Vehicle (VEH) 15 minutes before homecage behavior recording, 45 min before blood collection and 105 minutes before brain extraction. (b) quantification of grooming in seconds (t(13) = 3.089, p = 0.0086) and a detailed home-cage behavioral sequelae following VEH (top) or NESS (bottom) administration. (c) Circulating corticosterone (CORT) levels following VEH or NESS administration (t(13) = 2.318, p = 0.0374). (d) c-fos immunohistochemistry expression following VEH (left) or NESS (right). Scale bars: 150um *p = <0.05 **p = <0.01. (e) Quantification of c-fos density in the PVN following VEH or NESS (t(13) = 2.459, p = 0.0287).

To determine which eCB ligand mediates tonic suppression of stress-like responses, we utilized pharmacological inhibition of both AEA and 2-AG biosynthesis. AEA is primarily synthesized by NAPE-PLD and an inhibitor of this enzyme, LEI401, has recently been developed and produces robust reductions in central AEA content (Mock et al., 2020). Using this tool, we sought to determine if depletion of AEA produced alterations in homecage grooming behavior and HPA axis activity, similar to AM251 and stress exposure (Fig.3a). The NAPE-PLD inhibitor, LEI401, was administered systemically 2h before recording behaviors. Interestingly, LEI401 treated mice exhibited significantly more stereotypic grooming behaviors when compared to animals who received VEH injections t(13) = 5.535, p < 0.0001 (Fig 3c). Consistent with this, LEI401 administration also activated the HPA axis with marked increases in circulating CORT levels (Figure 3d; t(8) = 2.947, p = 0.0185) and c-fos expression in the PVN (Fig 3 e, f; t(8) = 2.694, p = 0.0273).

**Figure 3.**
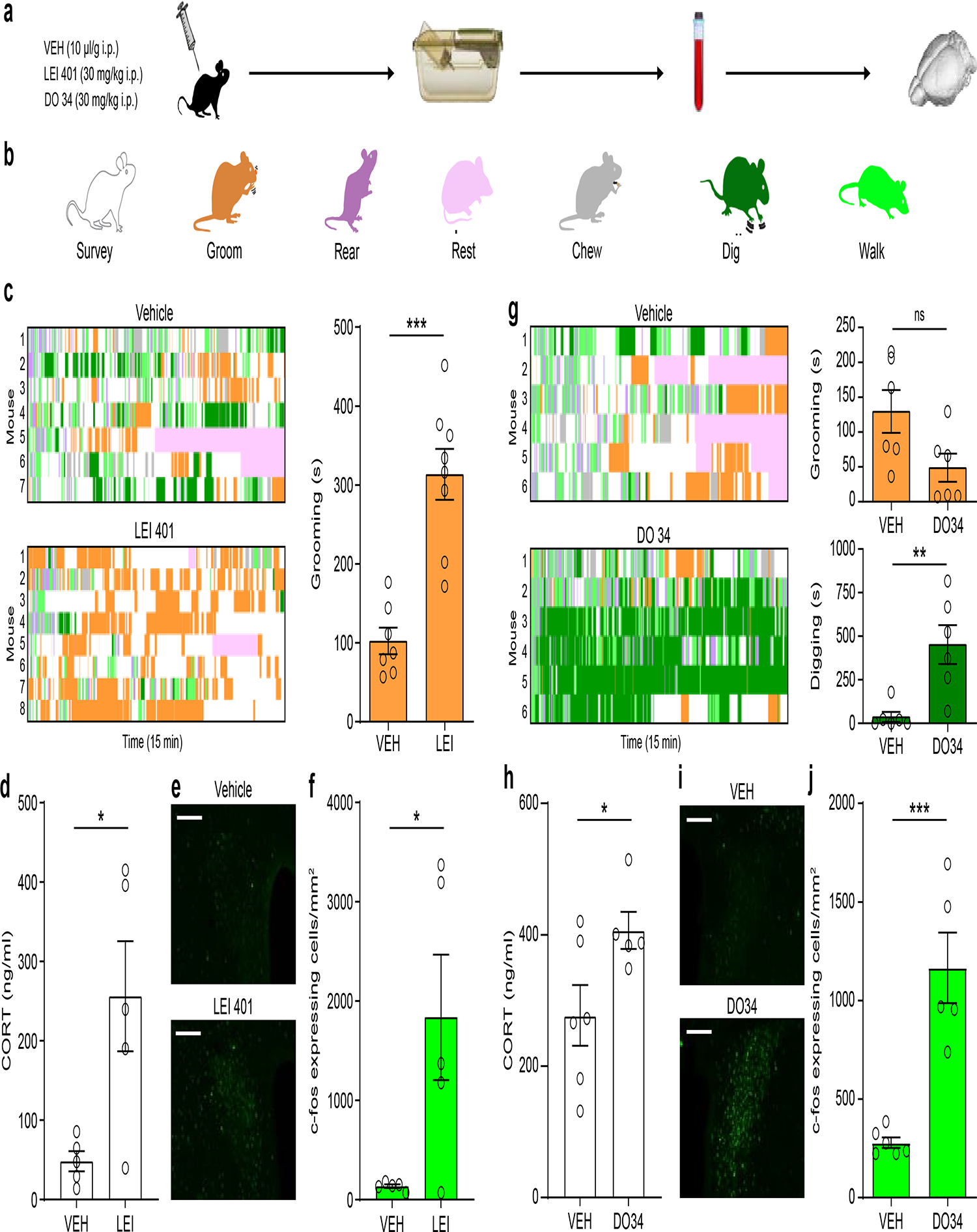
Inhibition of endocannabinoid biosynthesis activates a stress response. (a) LEI401 or DO34 were administered 2 h before home cage behavior recording, 2.5 h before blood collection and 3.5 h before brain extraction. (b) Color index of quantified behaviors throughout the behavioral epoch. (c) Detailed home-cage behavioral sequelae following VEH (top) or LEI401 (bottom) administration and quantification of grooming in seconds (s) (t(13) = 5.535, p < 0.0001). (d) Quantification of circulating CORT (t(8) = 2.947, p = 0.0185). (e) Representative images of c-fos immunohistochemistry expression following VEH (left) or LEI401 (right); Scale bars: 150um. (f) Quantification of c-fos in the PVN following VEH or LEI401 (t(8) = 2.694, p = 0.0273). (g) Detailed home-cage behavioral sequelae following VEH (top) or DO34 (bottom) administration and quantification of digging (t(10) = 3.602, p = 0.0048) and grooming (t(10) = 2.192, p = 0.0532) in seconds (s). (h) Circulating corticosterone (CORT) levels following VEH or DO34 administration (t(17) = 2.275, p = 0.049). (i) representative images of c-fos immunohistochemistry expression in the PVN; Scale bars: 150um. (j) Quantification of c-fos density in the PVN (t(9) = 5.402, p = 0.0004) and (following VEH (left) or DO34 (right) *p = <0.05 **p = <0.01 ***p = <0.001.

While many studies have suggested that AEA may be important for tonic eCB signaling (Hill et al., 2009; Gray et al., 2015; Natividad et al., 2017; Yasmin et al., 2020), there is also evidence that 2-AG may have tonic signaling capabilities (Lee et al., 2015; Marcus et al., 2020). To examine the impact of pharmacological depletion of 2-AG on stress-related measures, we inhibited 2-AG biosynthesis via administration of the DAGL inhibitor DO34 (Ogasawara et al., 2015). Two hours following DO administration, behavioral recordings revealed no differences in grooming behaviors when compared to VEH administration t(10) = 2.192, p > 0.05 (Fig. 3g). We further investigated the behavioral profile, which revealed that DO34 produced dramatic significant increases of repetitive digging behaviors t(10) = 3.602, p = 0.0048 (Fig. 3g), suggesting the possibility that the excess engagement in this behavior masked any notable differences in grooming behavior. However, animals receiving the DAGL inhibitor exhibited robust elevations in circulating CORT t(17) = 2.275, p = 0.049 (Fig. 3h), and c-fos expression in the PVN t(9) = 5.402, p = 0.0004 (Fig 3i,j), indicating an activation of the HPA axis and stress-like response. Overall, it appears that both AEA and 2-AG tonically regulate the HPA axis in a low stress environment, but the behavioral profiles evoked by these two treatments do differ, with depletion of AEA directly mirroring the effects of CB1 receptor antagonism or stress exposure (Fuzesi et al., 2016), while depletion of 2-AG produces large scale changes in different repetitive behaviors, but produces activation of neurons in the PVN and elevations in circulating CORT.

### The PVN is a critical hub for eCB regulation of the HPA axis

CRH neurons in the PVN orchestrate many aspects of the stress response, and previous work has suggested that the ability of stress to increase grooming behavior also requires activation of these neurons (Fuzesi et al., 2016). To determine if disruption of eCB signalling activated CRH neurons in the PVN to drive stress-like behavioral changes, we utilized optogenetics. Using the same technique employed in prior acute stress studies (Füzesi et al., 2016), the PVN of CRH-Cre mice was infected with a cre-dependent inhibitory opsin, Arch3.0- eYFP or a control eYFP virus (Fig. 4a). Following expression of the virus and implantation of ferrules over the PVN, mice were given either AM251 or VEH prior to behavioral recording. During home cage recording, continuous yellow light (532nm) was delivered over the PVN (Füzesi et al., 2016) (Fig.4b). One way ANOVA reported an effect of treatment (F(2, 15) = 9.425, p = 0.0022), and post hoc analysis revealed that within the group expressing eYFP, consistent with our initial study, AM251 significantly increased grooming behaviors in mice administered VEH (p=0.0012). In contrast, animals expressing Arch3.0 exhibited less AM251 induced grooming than animals expressing eYFP (p=0.0152) and did not differ from eYFP animals administered VEH (p=0.1971) (Fig. 4c,d). Thus, inhibiting CRH neurons in the PVN prevents AM251 from elevating stereotypic grooming similar to what has been established with stress.

**Figure 4.**
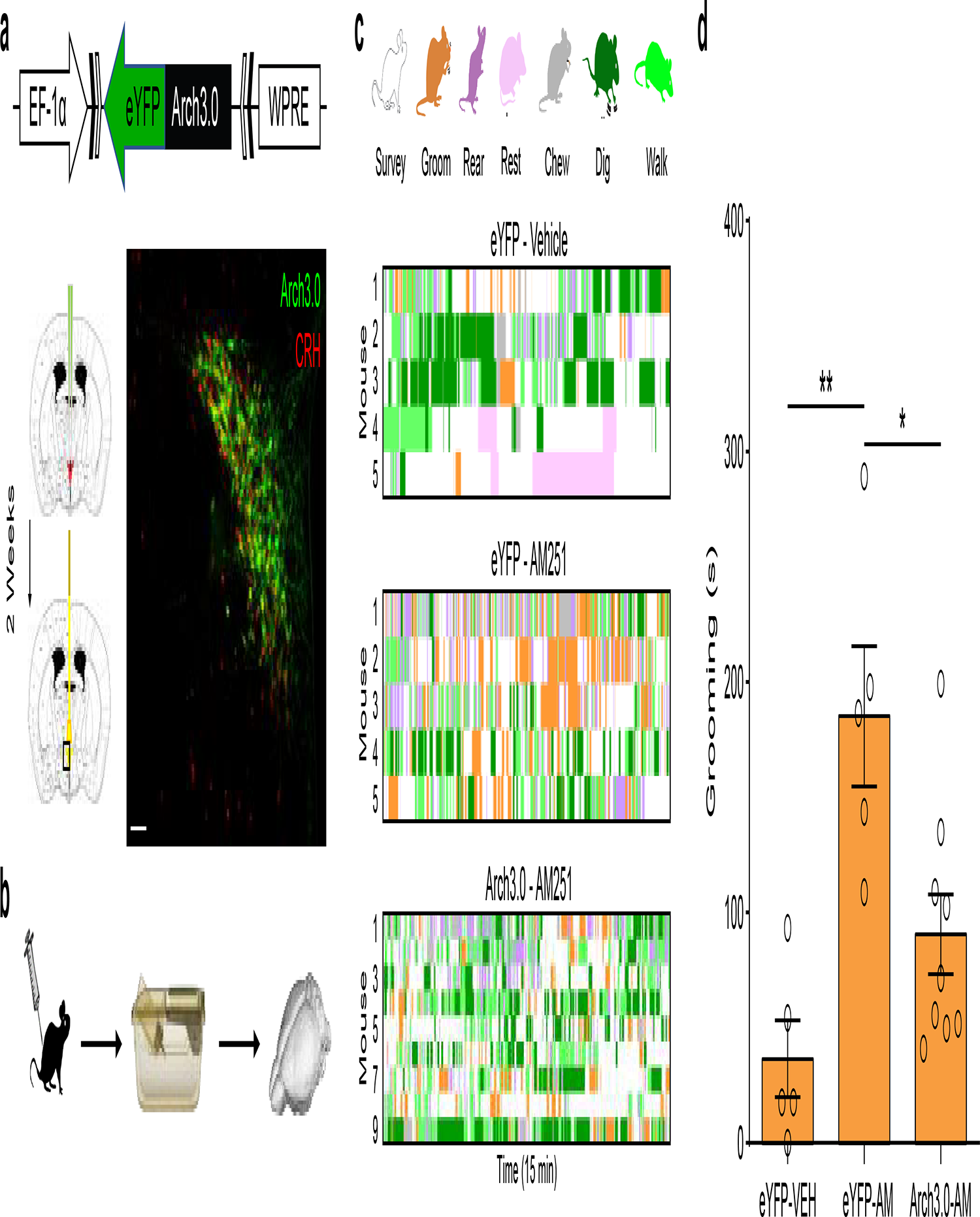
Inhibition of CRH neurons in the PVN blocks AM251 induced self-directed behaviors. (a) Cre-dependent AAV DIO Arch3.0-eYFP virus injected bilaterally into the PVN. (b) Animals were injected with VEH or AM251 15 minutes prior to constant 593nm light exposure inside the homecage. (c) Detailed behavioral output over the entire 15-minute behavioral epoch for each mouse. (d) Histogram analyzing grooming behavior in seconds (s) between groups. **p = <0.01.

Based on the optogenetic data provided above, local CRH neurons in the PVN appear to be orchestrating the stress-like stereotypic grooming following AM251 administration, similar to what is seen following acute stress. CB1 receptors are expressed within the PVN proper and activation of these receptors is known to locally influence excitatory and inhibitory synaptic transmission onto CRH neurons and regulate the stress response (Colmers and Bains, 2018; Di et al., 2003; Evanson et al., 2010; Nahar et al., 2015; Wamsteeker et al., 2010). As such, we next investigated whether local tonic eCB signaling in this region acts to gate the stress-like response. Following intracranial cannula implantation, AM251 was administered directly into the PVN 15 minutes before behavioral recording (Fig. 5a). Mice who received direct intra-PVN infusion of AM251 exhibited significantly increased stereotypic grooming behaviors when compared to VEH infused animals t(12) = 2.317, p = 0.0390 (Fig. 5d,e). Furthermore, intra-PVN AM251 infusions also resulted in elevated circulating CORT levels t(11) = 2.255, p = 0.0455 (Fig. 5f). These results suggest that tonic eCB signaling within the PVN proper is restricting activation of a stress response in ambient conditions and that disruption of this tone can produce the generation of a stress-like response. Based on the abundance of research pointing to the striatum as a regulator of grooming behaviors, we also investigated whether tonic eCB signaling was regulating self-directed behaviors here as well. Interestingly, administration of AM251 directly to the striatum had no effect of grooming behaviors t(13) = 1.231, p = 0.024 (Fig. 5h,i), indicating that this effect was at least partially specific to the PVN.

**Figure 5.**
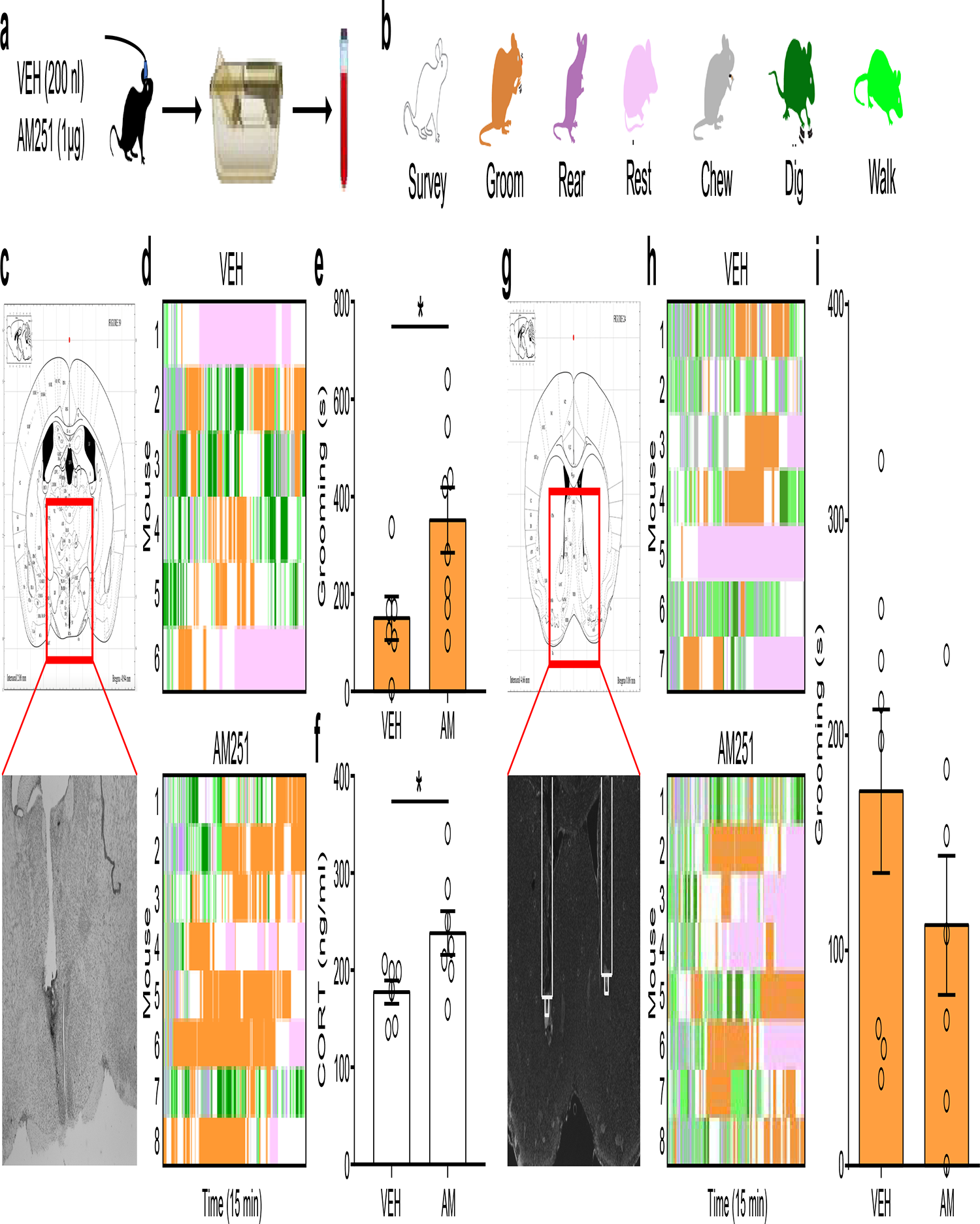
Local AM251 administration into the PVN initiates a stress-like phenotype. (a) Unilateral administration of VEH or AM251 15 minutes prior to homecage behavioral recording, 45 minutes before blood collection and 105 minutes before brain extraction. (b) Color index of quantified behaviors throughout the behavioral epoch. (c) histology of cannula placements into the PVN. (d) Detailed analysis of homecage behaviors following intra-PVN vehicle or AM251 over a 15-minute epoch and (e) histogram quantification of grooming behavior in seconds (s) following VEH or AM infusions (t(12) = 2.317, p = 0.0390). (f) Circulating corticosterone (CORT) levels following VEH or AM administration (t(11) = 2.255, p = 0.0455). (g) Bilateral administration of VEH or AM251 into the striatum 15 minutes prior to (h) homecage behavioral recording where (i) grooming was quantified (t(13) = 1.231, p = 0.24) alongside all other behaviors. *p = <0.05.

## Discussion

The current study expanded previous work regarding eCB regulation of the HPA axis and stress response by demonstrating that tonic signaling of both AEA and 2-AG, acting at the CB1 receptor within the PVN, restricts activation of the HPA axis at rest. Disruption of tonic eCB signaling produces rapid activation of the HPA axis and the manifestation of a behavioral stress-like response. Importantly, the manifestation of the stress-like response was driven by local actions of eCB signaling within the PVN proper and that optogenetic silencing of CRH neurons in the PVN inhibited this response. Together, these data indicate that tonic eCB signaling in the PVN restricts activation of CRH neurons in the absence of a threat or challenge, thereby providing an active mechanism of restraint on the HPA axis in ambient conditions.

AM251, as well as many other pharmacological tools used to block CB1 receptor signaling, are both antagonists and inverse agonists, making the mechanism by which they disrupt tonic eCB signaling complex. Disruption of eCB signaling could be due to blockade of eCB molecules binding to the CB1 receptor or it could be due to suppression of agonist-independent constitutive signaling of the CB1 receptor. Differential physiological and behavioral processes have been ascribed to both forms of tonic eCB signaling suggesting that both pathways are potential mechanisms (Lee et al., 2015; Meye et al., 2013; Sink et al., 2010). To address this question, we utilized a neutral antagonist, NESS0327, which only blocks agonist driven CB1 receptor activation but does not influence constitutive receptor signaling (Ruiu et al., 2003). Similar to what was found following administration of AM251, NESS0327 increased c-fos expression in the PVN, elevated CORT and increased grooming behavior. This indicates that tonic eCB regulation of CRH neurons in the PVN, and activation of the HPA axis, is driven by eCB molecules signaling directly on the CB1 receptor, and not constitutive CB1 receptor signaling.

Previous work has suggested that AEA is likely the eCB molecule mediating tonic inhibition of the HPA axis. AEA levels are known to decline in response to stress, and inhibition of AEA metabolism by fatty acid amid AEA hydrolase (FAAH) attenuates many endocrine and neurobehavioral responses to stress (Morena et al., 2016). The amygdala, particularly the basolateral nucleus (BLA), is a hub for some of the stress constraining effects of AEA (Bedse et al., 2014; Ganon-Elazar and Akirav, 2009; Hill et al., 2009; Gray et al., 2015). AEA provides tonic inhibition onto glutamatergic afferents into the BLA, and a loss of AEA signaling following stress exposure removes this tonic inhibition, resulting in increased excitatory drive onto BLA pyramidal neurons (Bedse et al., 2017; Yasmin et al., 2020). It is possible a similar mechanism is at play within the PVN as CB1 receptors have been found to regulate excitatory afferents impinging upon CRH neurons in the PVN (Di et al., 2003; Malcher-Lopes et al., 2006; Mazier et al., 2019; Wamsteeker et al., 2010). However, synaptic studies do not indicate that this input is subject to tonic inhibition by eCB signaling (Di et al., 2003; Wamsteeker et al., 2010) like it is in the BLA (Bedse et al., 2017; Yasmin et al., 2020). Alternatively, local CRH neuron microcircuits, which regulate GABAergic inputs onto parvocellular neurons in the PVN (Jiang et al., 2018), could represent a different circuit through which alterations in eCB signaling could be acting to modulate the stress response. This effect could also involve local CB1 signaling on astrocytes, as eCB signaling can regulate the release of gliotransmitters (Covelo et al., 2021; Navarrete et al., 2014), and CB1 signaling on astrocytes is known to be important for other stress-related processes such as fear learning (Martin-Fernandez et al., 2017). Perhaps disruption of eCB signaling onto, or from, neighbouring astrocytes could influence the excitability of CRH neurons, thus regulating HPA axis activity. Further work is required to understand the mechanism by which local eCB signaling gates activation of CRH PVN neurons.

As disruption of 2-AG produced similar neuroendocrine effects to that of antagonism of the CB1 receptor or depletion of AEA, it seems likely that there is tonic 2-AG signaling that is constraining HPA axis activity. The lack of effect of 2-AG depletion on grooming behavior could be due to this pharmacological approach triggering a dramatic elevation in a different stereotyped behavior which masked changes in grooming behavior. Animals treated with DO34 spent the overwhelming majority of their time burying their noses in the bedding then plowing it back and forth throughout the cage. Others have referred to this behavior as “inchworming” and it has been identified as a stereotyped behavior sometimes seen in rodent models of autism (Smith et al., 2014). It is unclear what is driving this behavior to occur, but given that 2-AG signaling in the striatum is known to regulate motor behavior (Lerner et al., 2010; Shonesy et al., 2013), it seems possible that depletion of 2-AG results in the engagement of a striatal circuit that triggers this behavioral stereotypy, which, in turn, overrides the expression of grooming behavior. Further work is required to understand the nature and significance of this behavior, but regardless, the neuroendocrine evidence seen herein indicates that 2-AG signaling does indeed provide tonic inhibitory control over the HPA axis.

While the canonical view is that CRH neurons are primarily neuroendocrine regulators, accumulating evidence over the past few years has demonstrated that this discrete cluster of neurons also has important and complex roles in coordinating behavioral responses to varying degrees of threat and reward (Daviu et al., 2020; Li et al., 2020; Sterley et al., 2018; Yuan et al., 2019). Interestingly, many of these processes are similarly influenced by changes in eCB signaling. As the ability of AM251 to trigger grooming responses was inhibited by optogenetic silencing of PVN CRH neurons, it will be important to understand if eCB modulation of other aspects of threat and defensive behavior will similarly involve modulation of PVN CRH neurons. Despite grooming being a relatively non-specific behavior that can be triggered by a host of stimuli and be mediated through different circuits, the current data do indicate that its expression is dependent on activation of PVN CRH neurons, similar to what is seen following stress (Füzesi et al., 2016). This is consistent with the fact that direct intra-cranial administration of CRH reliably triggers grooming behavior as well (Gargiufo and Donoso, 1996; Sherman and Kalin, 1987, 1986). Despite the well known importance of striatal circuits in driving grooming behaviors (Ahmari et al., 2013; Ramírez-Armenta et al., 2022; Zhang et al., 2021), consistent with our other data, local administration of AM251 into the PVN produced an increase in grooming, while a comparable manipulation into the striatum had no effect. That being said, a direct connection from CRH PVN neurons to the globus pallidus has recently been identified (Hunt et al., 2018), providing an anatomical substrate through which changes in CRH PVN neuron activity could rapidly modulate motor circuits to trigger grooming.

Taken together, these data clearly establish that tonic signaling of AEA and/or 2-AG, within the PVN, regulates the activation of CRH neurons in the PVN. Disruption of this tonic eCB signaling triggers the assembly of a neurobehavioral and endocrine phenotype that parallels what occurs in response to stress exposure. These data confirm that disruption of eCB signaling is sufficient to generate a stress-like response using multiple readouts and various pharmacological approaches. Both rodent and human studies have found that stress can produce declines in eCB signaling (Hill et al., 2009; Mayo et al., 2020a; McLaughlin et al., 2012; Patel et al., 2005; Rademacher et al., 2008; Spohrs et al., 2022; Yasmin et al., 2020), suggesting that a loss of eCB signaling may not only be sufficient, but also necessary, for the initiation of a stress-like response. A high proportion of humans who were taking the anti-obesity drug Rimonabant, which was a CB1 receptor antagonist, discontinued their use due to the onset of stress-related psychiatric symptoms, particularly anxiety and depression (Christensen et al., 2007; Hill and Gorzalka, 2009). These outcomes are consistent with the consequence of a persistently enhanced drive on PVN CRH neurons due to chronic disruption of eCB signaling. These data may also help to explain the low stress/anxiety phenotype seen in humans carrying a mutation in the FAAH gene, which results in reduced AEA metabolism and tonic elevations in eCB signaling (Dincheva et al., 2015; Gunduz-Cinar et al., 2013; Hariri et al., 2009; Mayo et al., 2020a; Spagnolo et al., 2016). Perhaps the basal elevations in eCB tone produced by this gene variant provide enhanced restriction of stress responsive neural circuits, centrally involving CRH neurons in the PVN. As FAAH inhibitors are currently under investigation for the management of anxiety and stress-related conditions (Mayo et al., 2020b; Paulus et al., 2021; Schmidt et al., 2020), with some encouraging data published to date from human clinical trials, this ability of tonic eCB signaling to constrain PVN CRH neuron activity may prove to be an important mechanism in this process. Collectively, these data highlight the importance of eCB signaling in the regulation of stress and the HPA axis and indicate that CRH neurons in the PVN are an important nexus for this interaction.

## Author Contributions

GNP and MNH designed the experiments, GNP performed the majority of experiments and wrote the initial draft of the manuscript and made the figures and worked with MNH on revising and finalizing the manuscript. GB, TF and RJA all made substantial contributions to the acquisition of data, revising the manuscript and assisting with figure development. WD and MvdS contributed to the acquisition of data by providing critical pharmacological agents and assisting with revising the manuscript. JSB contributed to the conception and design of the studies and provided substantial input on the revisions of the manuscript.

